# *FvFT1-FvTFL1* epistasis drives flowering time adaptation in woodland strawberry

**DOI:** 10.1101/2025.06.11.659068

**Authors:** Sergei Lembinen, Elli Koskela, Guangxun Fan, Tuomas Toivainen, Quan Zhou, Javier Andrés, Aleksia Vaattovaara, Pasi Rastas, Hanne De Kort, Paula Elomaa, Timo Hytönen

## Abstract

Floral transition and flower bud development during autumn and winter are common in plants inhabiting temperate and cold climates. However, our understanding of the natural variation and underlying molecular mechanisms of such life-history strategy remain largely limited to Brassicaceae model species. To address this gap, we investigated flowering time variation across a broad geographical and bioclimatic range in *Fragaria vesca* (woodland strawberry), a perennial herb from the Rosaceae family. We found that the timing of floral transition and subsequent flowering in *F. vesca* populations is primarily shaped by population structure, reflecting postglacial colonization patterns from distinct refugia, and is further modulated by adaptations to winter temperatures. The association of increasing expression of *FvFT1*, the strawberry ortholog of *FLOWERING LOCUS T*, with delayed flowering in accessions from diverse climatic and genetic backgrounds suggested *FvFT1* as a major candidate gene regulating flowering time variation. We further demonstrated how distinct florigen and antiflorigen roles of FvFT1 are dependent on functional alleles of *F. vesca* TERMINAL FLOWER1 (FvTFL1) and how the epistatic relationship between *FvFT1* and *FvTFL1* controls adaptive flowering time variation by adjusting sensitivity to floral inductive photoperiod and vernalization. Our findings reveal a major molecular basis of how timing of floral transition has adapted to temperature in a Rosaceae model inhabiting multiple climatic zones and highlight lineage-specific adaptive mechanisms underlying convergent life-history strategy.

**Significance statement:** Molecular mechanisms underlying flowering time adaptation have been primarily investigated in annual plants that transit to flowering in spring. However, many perennial plant species in temperate climates form flower buds in autumn and winter to flower in the following spring. We uncover wide geographical and climatic patterns of flowering time adaptation in the perennial model species woodland strawberry. We report that natural variation in the expression of strawberry *FLOWERING LOCUS T* (*FvFT1*) gene originating ice ages ago and further driven by local temperature adaptations control flowering time variation in this species. Finally, we show how *FvFT1* may function as both flowering promoter and repressor depending on the genetic background, thus revealing a novel mechanism of flowering time adaptation.

## Introduction

Successful reproduction of plants requires precise timing of flowering aligned with the growing season. In many annual species that complete their life cycle in a single growing season, floral transition and initiation of floral primordia occur in spring and summer in response to increasing daylength and temperature. Molecular mechanisms controlling floral transition in these summer annuals typically converge to regulate the expression of a small signaling peptide, the FLOWERING LOCUS T (FT; Andrés and Coupland 2012). Natural variation at the *FT* locus and its upstream regulators is strongly associated with geographical flowering time variation allowing the plants to adapt to their seasonal environments (Schwartz et al. 2009, Liu et al. 2014, Rosas et al. 2014, Strange et al. 2011). In temperate climates, however, a common life-history of perennials and winter annuals comprises of formation of overwintering floral buds in autumn and flowering in the following spring (Schnablová et al. 2020). Floral transition and initiation of floral buds in such plants occur in cool temperatures and short days, shortly after a period of vernalization varying from weeks to months (O’Neill et al. 2019, Soppe et al. 2021). A more advanced developmental stage of floral buds before winter enables earlier flowering upon the onset of favorable conditions in spring which is essential in the environments with short growing season (Molau et al. 2005, Schnablová et al. 2020, Mauracher & Wagner 2021). Despite the wide distribution of this life-history strategy in temperate climates, our understanding of geographical patterns of flowering time variation along with the underlying molecular mechanisms and their adaptive natural variation is limited to a few models in the Brassicaceae family.

In winter annual and perennial Brassicaceae, orthologs of a MADS-box transcription factor FLOWERING LOCUS C (FLC) function as strong floral repressors, and a period of cool temperatures called vernalization is needed to silence *FLC*, enabling floral transition. For example, in perennial *Arabis alpina*, silencing of *FLC* in the shoot apical meristem during exposure to cold enables the activation of flowering promoter *SUPPRESSOR OF OVEREXPRESSION OF CONSTANS1* (*SOC1*) and floral meristem identity genes *LEAFY* (*LFY*) and *APETALA1* (*AP1*) leading to floral transition in late autumn/early winter (Wang et al. 2009, Wang et al. 2011, Lazaro et al. 2019). Furthermore, in winter annual and perennial Brassicaceae, natural variation in *FLC* expression levels and downregulation rates during vernalization contributes to the variation in the temperature requirement and duration of vernalization needed for floral transition. These differences in sensitivity of vernalization underlie flowering time variation across wide geographical ranges (Albani et al. 2012, Kemi et al. 2013, Duncan et al. 2015, O’Neill et al. 2019, Hepworth et al. 2020, Calderwood et al. 2021, Wunder et al. 2023, Zhu et al. 2023). Beyond Brassicaceae, however, the role of *FLC* genes in regulation of vernalization and floral transition appears to be less conserved, and other floral repressors need to be silenced during autumn or winter (Bouché et al. 2017).

Our studies are focusing on the regulation of flowering in woodland strawberry (*Fragaria vesca,* Rosaceae), a perennial herb widely distributed in the northern hemisphere (Hilmarsson et al. 2017). Floral transition in woodland strawberry occurs during shortening days and lowering temperatures of autumn, and overwintered flower buds open in the following spring. In this species, a homolog of TERMINAL FLOWER1 (FvTFL1) has replaced FLC as a major repressor of floral transition (Koskela et al. 2012). Similarly to *FLC* in *A. alpina* (Wang et al. 2009), decreasing temperatures gradually downregulate *FvTFL1* expression in the shoot apical meristem. However, the temperature limits of the vernalization response in woodland strawberry depend on photoperiod; in short days, temperatures below ∼18°C silence *FvTFL1* and induce floral transition, while at temperatures lower than 13°C *FvTFL1* is downregulated independently of photoperiod (Koskela et al. 2012, Rantanen et al. 2015). In *fvtfl1* mutants, in contrast, long-days and warm temperatures accelerate floral transition by inducing the expression of strawberry *FT* ortholog (*FvFT1*), specifically in the leaves, which promotes the expression of strawberry *LFY* and *AP1* homologs in the apical meristems (Koskela et al. 2012, Lembinen et al. 2023).

Recent findings demonstrated that European woodland strawberry populations are divided into eastern and western genetic clusters formed by postglacial northward colonization patterns from distinct refugia (Toivainen et al. 2024). These populations span a wide range of environments and likely harbor diverse hallmarks of flowering adaptation. For example, natural variation at the *FvTFL1* locus caused an extended vernalization requirement at low temperatures in an arctic subpopulation of *F. vesca* (Koskela et al. 2017). Here, we report large-scale geographical patterns of flowering time variation in European woodland strawberry. Our data highlights winter temperature as a major driver of flowering time adaptation in this species. Furthermore, we identify a link between flowering time variation and the expression level of *FvFT1* prior to floral transition. We further demonstrate a novel mechanism by which *FvFT1* controls sensitivity to floral inductive conditions (e.g. vernalization and short days) via activation of *FvTFL1*. Finally, we show how *FvFT1/FvTFL1* epistasis confers the dual role of *FvFT1* as a florigen and antiflorigen of strawberry.

## Results

### Patterns of flowering time variation in European woodland strawberry

To understand patterns of flowering time variation in European woodland strawberry, we subjected a collection of 179-189 accessions to flower inductive conditions in the field (Helsinki, Finland) and controlled climate followed by flowering time observations in a greenhouse (LD 20°C) or in the field. After exposing the plants to field conditions for different durations (Aug-Oct and Aug-Jan), all tested accessions flowered when moved to the greenhouse. Similarly, all accessions flowered when grown in the field during autumn and the following spring and summer (Sep-Jun). Almost all accessions flowered after short-day (SD 16°C, 174/179), cool temperature (LD 11°C, 175/182) or artificial seasonal cycle (SD 11°C/6°C/-2°C, 188/189) treatments in controlled climate, demonstrating that short days and cool temperatures in autumn largely behave as interchangeable flower-inductive signals for most *F. vesca* accessions across Europe. Flowering time strongly correlated between all tested experimental conditions (**Supplemental Figure S1A**) indicating that the rate of floral transition is the major determinant of flowering time in woodland strawberry. However, exposure to late autumn conditions in the field (Aug-Jan, Sep-Jun) or controlled climate (6°C) and overwintering (−2°C) decreased flowering time variation (**Supplemental Figure S1B**) by accelerating flowering in the accessions which flowered late in the other treatments. This suggests that some accessions may continue floral development later during autumn.

We further analyzed the geographical and bioclimate-related (Fick and Hijmans 2017) patterns of flowering time variation. In most experiments, flowering time correlated strongly with latitude, longitude, and temperature-related bioclimate variables characteristic of the habitat of the accession (**Figure 1A**). The strongest correlations were observed between flowering time and the variables representing winter temperature (e.g., bio6 and bio11) and seasonal temperature variability (e.g., bio3 and bio4), whereas correlations between flowering time and summer temperatures were weaker (e.g., bio5 and bio10) (**Supplemental Figure S1C**). Overall, the accessions originating from the habitats with colder, more seasonal climates typically flowered earlier, following both south to north and coast to inland clines (**Figure 1B**). However, the relationships between flowering time, latitude, and temperature were not linear (**Supplemental Table S1**). The latitudinal cline in flowering time was mild from latitudes of 37°N to 60°N, while at latitudes above 60°N the cline decreased rapidly (**Figure 1C, Supplemental Figure S2**). Furthermore, for winter temperature (bio11), a strong positive cline was only visible for the accessions originating from colder habitats, representing bio11 values from –12°C to 0°C, while above 0°C, the cline flattened (**Figure 1D**). These changes in the patterns of flowering time variation for latitude and winter temperature were largely associated with previously reported eastern and western genetic clusters of European woodland strawberry (**Figure 1B – D**; Toivainen et al. 2024). The accessions belonging to the western genetic cluster are typically found in warmer climates and flowered later in all experimental treatments (**Supplemental Figures S3-4**). Linear regression analysis confirmed steeper clines for latitude and bio11 variables in the eastern cluster in all experimental treatments (**Figure 1E, Supplemental Table S2**). Furthermore, stronger clines were found for bio11 than latitude suggesting that temperature rather than photoperiod has a dominant role in flowering adaptation in European *F. vesca*.

**Figure 1.**
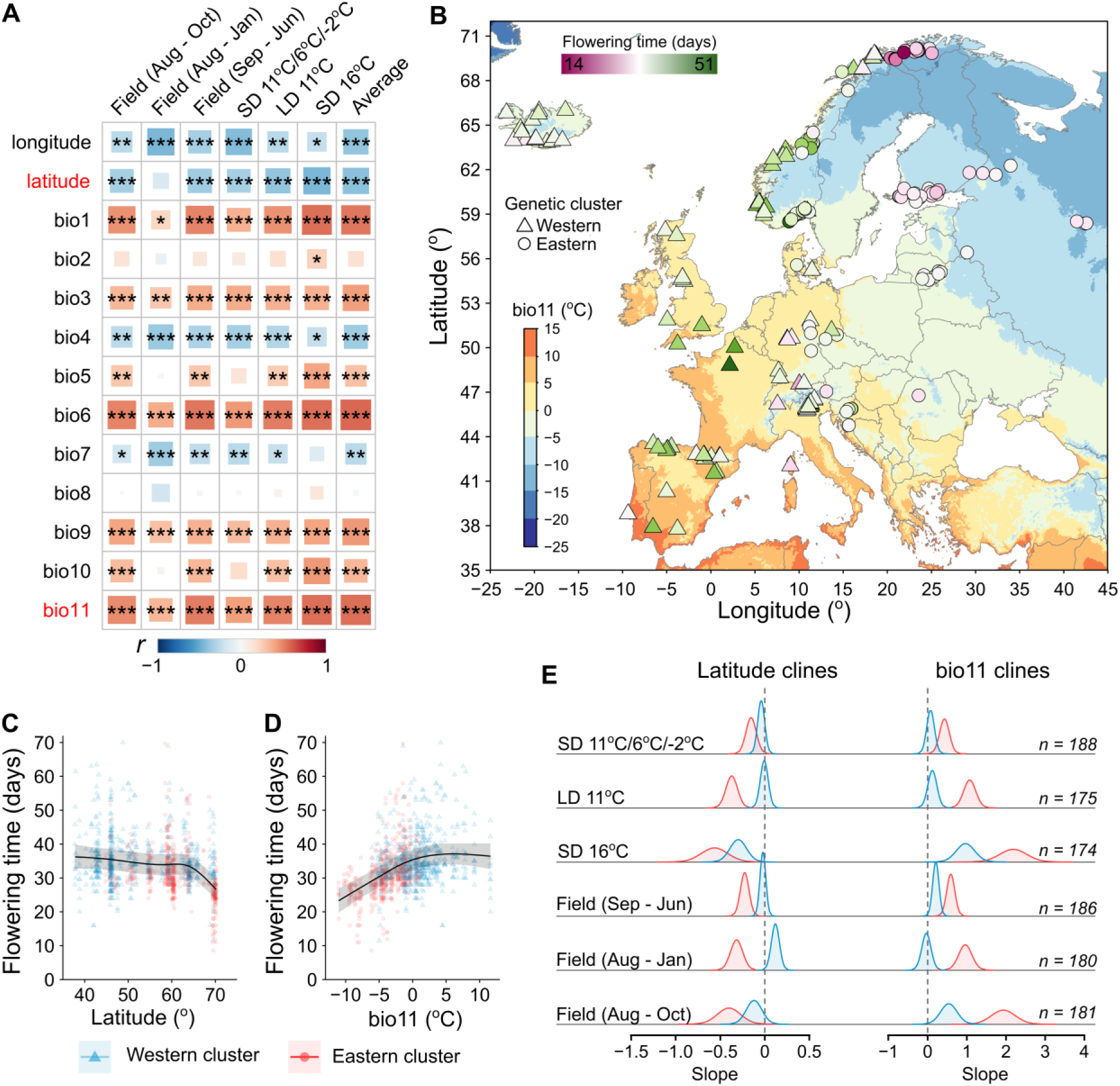
Patterns of flowering time variation in European woodland strawberry populations. **A.** Heatmap showing correlations (Pearson’s) between flowering time in different experiments and temperature related bioclimatic variables, longitude and latitude. Asterisks mark the significance levels (***, p < 0.001; **, p < 0.01; *, p < 0.05, Bonferroni adjusted). **B.** Map showing collection sites of woodland strawberry accessions and mean temperature of the coldest quarter (bio11). Color of symbols indicates average flowering time from six different experiments and their shape denotes genetic clusters. **C** – **D.** Non-linear patterns of flowering time variation along latitude (**C**) and temperature (bio11) (**D**) gradients. **E.** Estimates of latitudinal and bio11 slopes of flowering time variation for eastern and western genetic clusters in six different experiments.

The effects of population structure and winter temperature on flowering time were dominant even at extremely high latitudes. This is supported by flowering time observations in clonal samples gathered from northern Norway, a region with closely located but highly diversified populations (Tromsø, Kåfjord and Alta) and habitat bioclimates (Toivainen et al. 2024; **Supplemental Figure S5A – B**). The populations of Tromsø region occupy warmer habitats and belong to the western genetic cluster, whereas the near-by populations of Alta and Kåfjord belong to the eastern cluster and occupy colder habitats. In both controlled climate and common garden experiments, the clones from the colder habitats of Alta and Kåfjord flowered earlier than the clones from the warmer habitats of Tromsø (**Supplemental Figure S5C**). Taken together, our data suggests that postglacial northward colonization patterns from distinct refugia and further adaptation to winter temperature have been the main drivers in shaping the current patterns of flowering time variation in European woodland strawberry.

### Cis- and trans-regulation of *FvFT1* contribute to adaptive flowering time variation in woodland strawberry

We next aimed at identifying candidate gene(s) controlling flowering time variation in accessions across Europe. GWAS using whole-genome data was unsuccessful in finding stable significant markers associated with flowering time, likely due to complex population structure consisting of isolated or semi-isolated highly differentiated populations (Toivainen et al., 2024). As an alternative approach, we used RNA sequencing of leaf tissue obtained from 12 selected accessions from the eastern and western genetic clusters spanning the temperature range of woodland strawberry distribution (**Supplemental Data S2**, **Supplemental Figure S6A**). Among the 659 differentially expressed genes between the pooled eastern and western accessions, only three flowering-related genes were identified, namely *FvFT1*, *FvTPS1* (*TREHALOSE-6-PHOSPHATE SYNTHASE1*) and *FvLUX* (*LUX ARRHYTHMO*) (**Figure 2A**). Both *LUX* and *TPS1* were previously identified as upstream regulators of *FT* expression (Hazen et al., 2005, Wahl et al., 2013). We therefore reasoned *FvFT1* as the most relevant candidate to control wide-scale flowering time variation in European *F. vesca*. To test this hypothesis, we analyzed *FvFT1* expression in a larger set of 48 accessions, spanning the geographical range of our collection (**Supplemental Data S1**). The increasing relative expression level of *FvFT1* was strongly associated with delayed flowering (r = 0.47, p = 8.19 × 10^-4^), higher bio11 values (r = 0.73, p = 5.47 × 10^-9^), and lower latitude (r = –0.61, p = 3.44 × 10^-6^) (**Figures 2B – D**), suggesting a role for *FvFT1* in repression of flowering and a link between *FvFT1* expression and the adaptive flowering time variation in European woodland strawberry.

**Figure 2.**
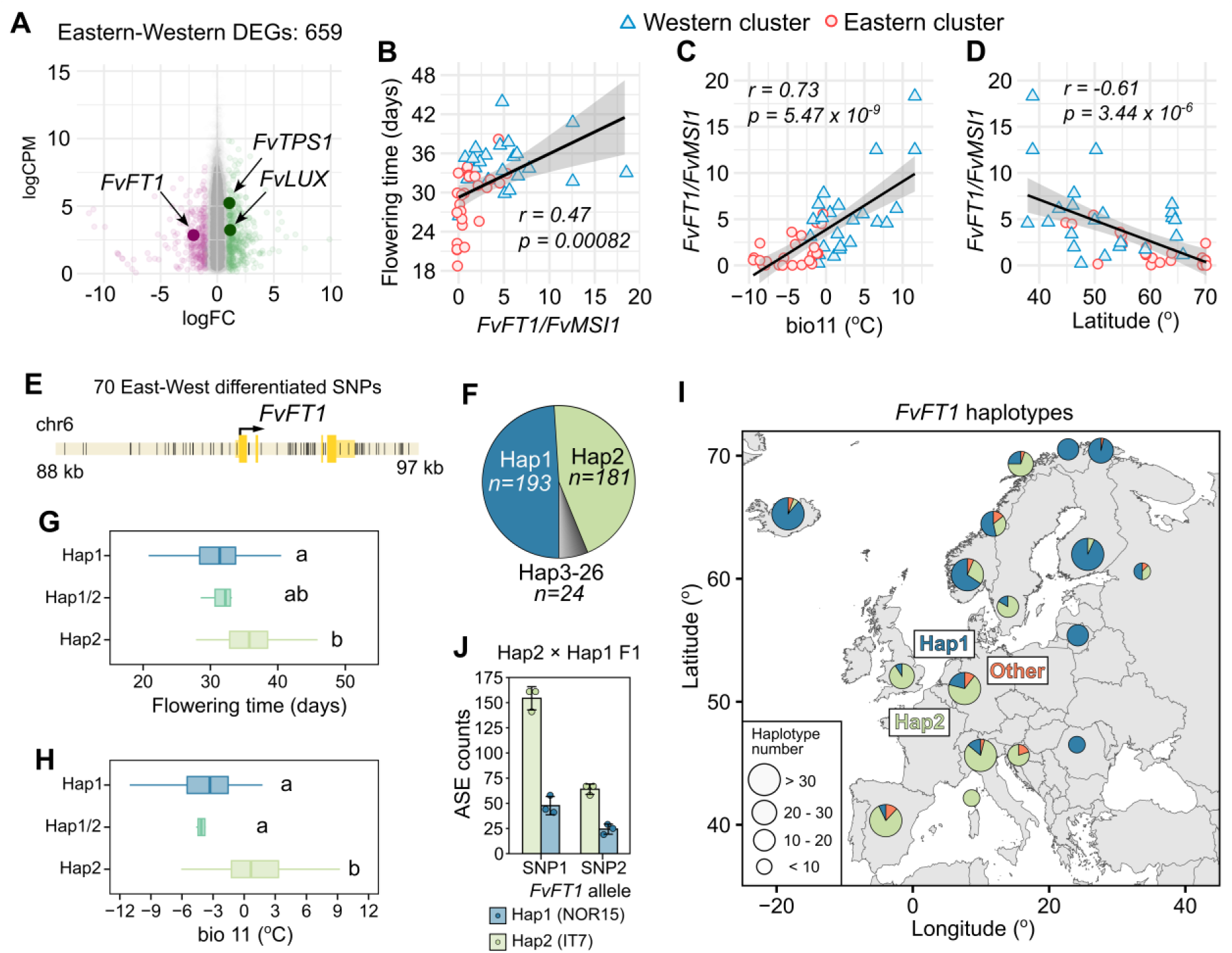
*Cis* regulation of *FvFT1* contributes to flowering time variation in European woodland strawberry. **A.** Differentially expressed genes (DEGs) between six western and six eastern accessions. **B – D.** Correlations (Pearson) between relative *FvFT1* expression level and flowering time (**B**), mean temperature of the coldest quarter (bio11) (**C**), and latitude (**D**). Genetic clusters are highlighted in different colors (Toivainen et al. 2024). **E.** Highly differentiated SNPs between the eastern and the western genetic clusters (among top 1% F_ST_) in *FvFT1* locus located within indicated position on the chromosome 6. **F.** Haplotype frequencies. **G.** Flowering time of the accessions carrying the major haplotypes. **H.** Comparison of winter temperatures (bio11) at the haplotype locations. Letters indicate significantly different groups (p < 0.05; Tukey’s HSD). **I.** Geographical distribution of three major *FvFT1* haplotypes. **J.** Comparison of *FvFT1* allele specific expression from Hap2 × Hap1 F_1_ cross.

To investigate the functional role of *FvFT1* in driving flowering time variation, we explored the genomic variation at *FvFT1* locus in the collection of 202 woodland strawberry accessions. We identified 70 highly differentiated SNPs between the eastern and western genetic clusters (belonging to the top 1% F_ST_ outliers across the genome) spanning the region of almost 9 kb and located in both 5’ (n = 25) and 3’ (n = 22) regulatory regions and introns (n = 23) (**Figure 2E; Supplemental Data S3**). A closer analysis revealed two major haplotypes of *FvFT1* locus (Hap1 and Hap2), and 24 minor haplotypes primarily found as single copies (**Figure 2F**). Hap1 was more abundant among the accessions of the early flowering eastern cluster, whereas Hap2 was typical for the western cluster (**Figure 2G**). An exception were the accessions from Iceland that belong to the western genetic cluster (Toivainen et al. 2024) but almost exclusively contain the eastern Hap1. The distribution of the haplotypes correlated with the bio11 values (**Figure 2H and I**). These results suggest a major role for *FvFT1* in temperature adaptation of flowering in woodland strawberry. To corroborate these findings, we analyzed the allele specific expression of *FvFT1* in F_1_ crossing progeny between the Italian and Norwegian accessions IT7 (Hap2) × NOR15 (Hap1) (**Supplemental Data S4**). Transcripts with two SNPs in *FvFT1* coding sequence were detected. Based on the read counts of each SNP, the alleles driven by Hap2 regulatory sequences were more abundantly expressed than the alleles of Hap1 (**Figure 2J**), consistent with differential *FvFT1* expression between eastern and western populations. Altogether, our data suggests that the geographic pattern of *cis*- regulatory variation in *FvFT1*, which likely began to develop several Ice Ages ago (Toivainen et al. 2024), contributes to flowering time variation.

To investigate the possibility of *trans*-regulation of *FvFT1*, we performed weighted gene co-expression network analysis (WGCNA; Langfelder & Horvath 2008) on the RNAseq data from 12 accessions and identified gene clusters highly correlated with flowering time, bio11, latitude and specific accession (**Supplemental Figure S6B - D, Supplemental Data S2**). Clusters 10-12, showing the strongest positive correlations with flowering time (**Figure 3A**), contained the strawberry homologs of several flowering-related genes including *FvFT1, SQUAMOSA PROMOTER BINDING PROTEIN-LIKE 3* (*FvSPL3*), and *FRUITFUL* (*FvFULa*) (**Figure 3B and C**). The expression differences of these genes were associated with the eastern and western genetic clusters. Furthermore, *FvTPS1* was found in cluster 4 that negatively correlated with flowering time in multiple experiments (**Figure 3A and B**). In addition, in some clusters, flowering genes putatively involved in *FvFT1* regulation showed population specific expression patterns, particularly in the accessions from the two early flowering populations from the adjacent fjords in northern Norway (**Figure 3A, D, E**). For example, in the accessions from Kåfjord (NOR14 and NOR15), the expression of strawberry orthologs of *CONSTANS (CO)*, *SOC1* and two *CYCLING DOF FACTOR2* (*CDF2*) genes were downregulated (cl10; **Figure 3D**). FvCO and FvSOC1 have previously been shown to function in photoperiodic regulation of flowering in woodland strawberry (Mouhu et al. 2013; Kurokura et al. 2017), and CDF2 is involved in the photoperiodic flowering in Arabidopsis (Fornara et al. 2009). In Alta accessions (NOR5 and NOR8), the expression of four *FRIGIDA-like* (*FRI-like*) genes closely located on the chromosome 6 was strongly downregulated or completely undetected (cl13; **Figure 3E**). This indicates that besides large-scale cis-regulatory variation in *FvFT1*, local populations evolved specific adaptations which may converge to regulate *FvFT1* expression.

**Figure 3.**
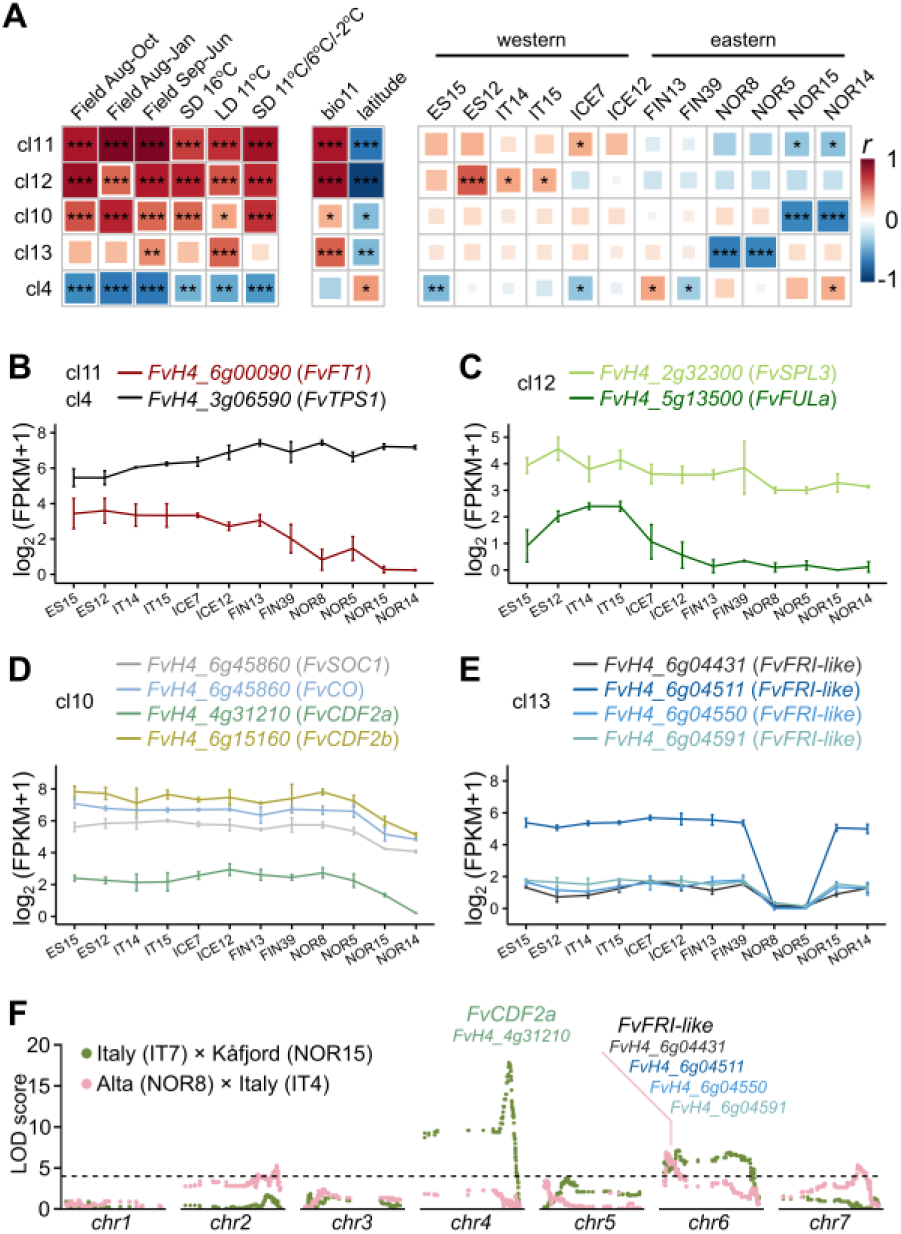
*Trans* regulation of *FvFT1* contributes to flowering time adaptation in local populations. **A.** Heatmap showing Pearson’s correlation between selected RNAseq clusters (WGCNA), flowering time in different experiments, mean temperature of the coldest quarter (bio11), latitude, and accessions. **B – E.** Expression patterns of selected genes from different clusters. **F.** LOD scores of markers from two (F_2_) mapping populations with candidate genes highlighted.

To further investigate population specific differences in flowering time regulation, we produced two small mapping populations using accessions from either Alta (NOR8) or Kåfjord (NOR15) as one of the parents. In IT7 × NOR15 F_2_ population, *FvCDF2a* was found close to the QTL peak for the flowering time (**Figure 3F; Supplemental Table S3**, **Supplemental Figure S7**). Furthermore, the locus containing *FvFRI-like* genes was within the confidence interval of the flowering time QTL in NOR8 × IT4 F_2_ population (**Figure 3F**). Although further functional validation is required, the different QTL peaks from the accessions from adjacent fjords suggest population specific mechanisms in the regulation of *FvFT1* expression and flowering time in European woodland strawberry.

### *FvFT1* controls sensitivity to flower inductive conditions

To further understand the role of FvFT1 in the regulation of flowering time variation, we first analyzed the expression of *FvFT1* and *FvTFL1*, the latter encoding a major flowering repressor (Koskela et al. 2012), in 40 accessions grown in non-inductive long days (**Supplemental Data S1**). The expression of *FvTFL1* in the shoot apex and *FvFT1* in the leaves were strongly positively correlated (r = 0.71, p = 2.42 × 10^-7^, **Figure 4A**), suggesting that the leaf expressed *FvFT1* positively control the pre-inductive levels of *FvTFL1* in the shoot apical meristem, and consequently, the flowering time. We used the accessions from Italy and northern Norway (Alta/Kåfjord) to test this potential mechanism. Accessions from northern Norway flowered earlier than the accessions from Italy (p = 5.0 × 10^-9^; **Figure 4B**) and had lower expression of *FvFT1* in their leaves (**Figure 4C**). Furthermore, the temporal dynamics of *FvTFL1* and *FvAP1* in the shoot apices under flower-inductive conditions were different between the accessions. Before the start of the inductive treatment (W0), the basal *FvTFL1* expression levels in Italian accessions were higher (W0, p = 0.023; **Figure 4D**), and remained higher until the end of the sampling period (W4, p = 0.018) although following a similar downregulation dynamic as in the Norwegian accessions. Concordantly, in the Norwegian accessions, upregulation of *FvAP1* (as indicator of floral transition) was observed already during the third week of the treatment (W3, p = 0.0038), after downregulation of *FvTFL1* under a critical threshold, whereas in the Italian accessions, the weak upregulation of *FvAP1* was detected only at week 4 (W4, p = 0.00032; **Figure 4E**). Altogether this suggests that the timing of floral transition is controlled by the initial expression level of *FvTFL1*, which is regulated by *FvFT1*.

**Figure 4.**
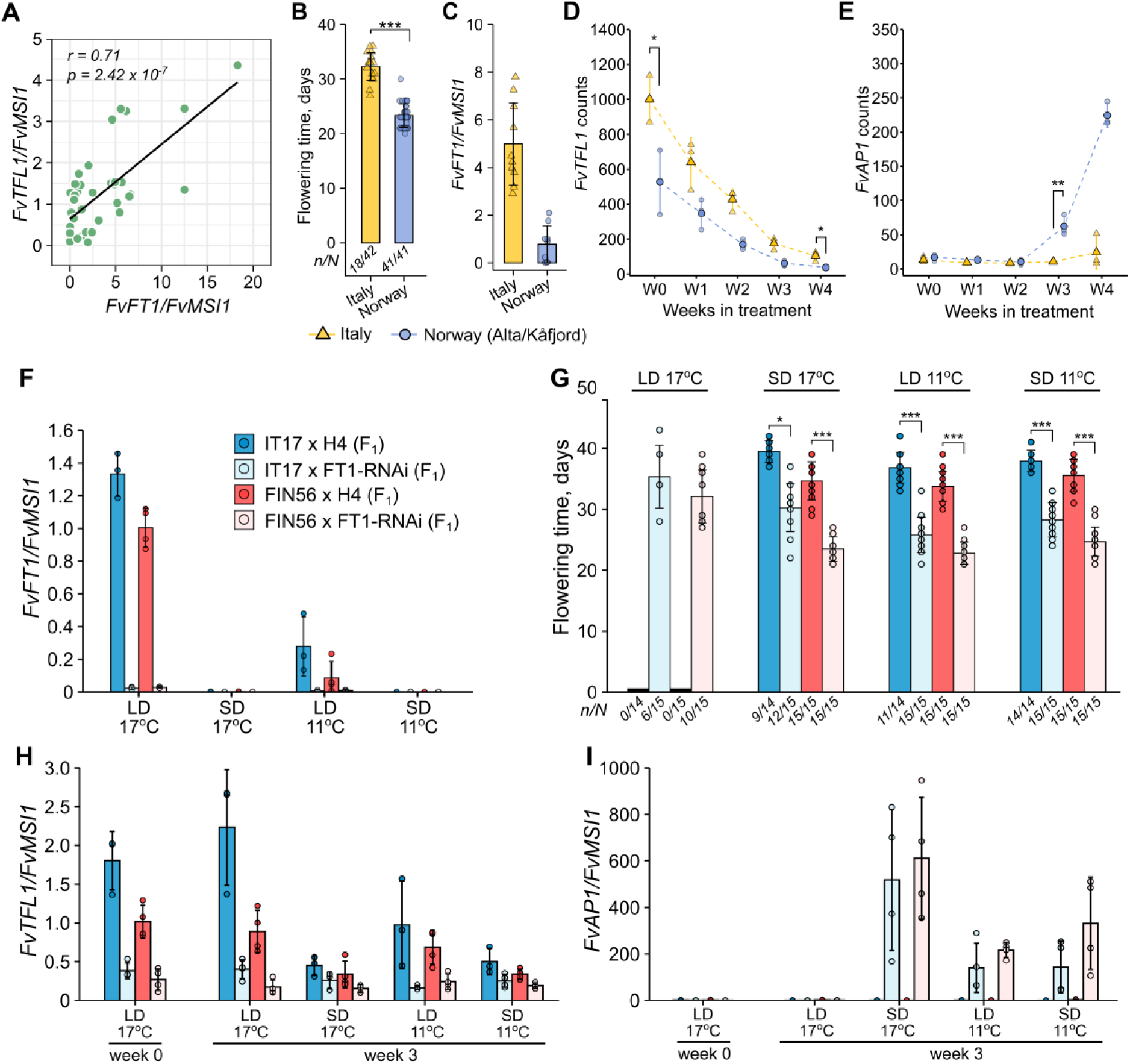
*FvFT1* controls sensitivity to flower inductive cues in woodland strawberry. **A.** Pearson’s correlation between relative expression of *FvFT1* in leaves (ZT4) and *FvTFL1* in shoot apices under non-inductive LD 17°C conditions in 40 different woodland strawberry accessions. **B.** Flowering time of woodland strawberry accessions from Italy and northern Norway (Kåfjord and Alta) after four weeks in LD 11°C treatment. Mean data of three Italian (IT4, IT12, IT17) and three Norwegian (NOR2, NOR8, NOR16) accessions with 11-15 replicates per accession is shown. Numbers above x-axis denote the number of flowered plants (*n*) and total number of plants in each group (*N*). **C.** Relative expression of *FvFT1* in leaves (ZT4) of Italian and northern Norwegian accessions under LD 17°C. **D – E.** Expression dynamics of *FvTFL1* (**D**) and *FvAP1* (**E**) in shoot apices over four weeks of LD 11°C treatment in accessions from Italy and northern Norway. Asterisks denote significant differences between groups at selected time points (***, p < 0.001; **, p < 0.01; *, p < 0.05; t-test). **F.** Relative expression of *FvFT1* in the leaves of transgenic *FvFT1-RNAi* and control crosses after three weeks under the indicated treatments. **G.** Flowering time (days after treatments) of woodland strawberry *FvFT1-RNAi* and control crosses after six weeks under the indicated treatments. y-axis shows days since the end of the treatments; asterisks indicate the significant differences between selected groups (***, p < 0.001; **, p < 0.01; *, p < 0.05; GLM (Poisson) with Tukey’s HSD test). Numbers under the x-axis denote the numbers of flowered plants (*n*) and the total number of plants in each group (*N*). **H – I.** Relative expression of *FvTFL1* (**H**) and *FvAP1* (**I**) in the shoot apices of transgenic *FvFT1-RNAi* and control crosses before and after three weeks of the treatments.

To further test the function of *FvFT1*, we crossed IT17 (western) and FIN56 (eastern) accessions with *FvFT1-*RNAi line in Hawaii-4 (H4) background (Koskela et al. 2012) and with H4 wild type (control). Because H4 is a *fvtfl1* mutant (Koskela et al. 2012), all F_1_ lines contained a single functional *FvTFL1* allele. The introduced *FvFT1*-RNAi construct effectively silenced *FvFT1* in both IT17 and FIN56 (**Figure 4F**). In addition, both flower-inductive short days and cool temperatures downregulated *FvFT1* in control plants. The RNAi silencing of *FvFT1* significantly shortened time to flowering under all experimental treatments (**Figure 4G**). In addition, 40% of IT17 and 67% of FIN56 *FvFT1*-RNAi plants were able to flower under non-inductive LD 17°C treatment supporting the role of *FvFT1* in repression of flowering. We then analyzed the expression patterns of *FvTFL1* and *FvAP1* in the shoot apex samples (**Figure 4H and I**). The expression of *FvTFL1* was lower under long-days and warm temperature in the plants with silenced *FvFT1*. After three weeks under short-days or under cool temperature in short or long days, *FvTFL1* was downregulated in both control and *FvFT1-*RNAi plants. However, even after three weeks of the treatments, the expression level of *FvTFL1* in control plants was higher than in the plants with silenced *FvFT1* prior to the start of the inductive treatments (week 0). Concordantly, only *FvFT1-*RNAi plants showed upregulation of *FvAP1* after three weeks of treatments indicating earlier flower induction. The results uncover a mechanism controlling sensitivity to flower inductive conditions in woodland strawberry. By adjusting the expression levels of *FvFT1* under non-inductive long days, plants can regulate the expression of flowering repressor *FvTFL1*, which is gradually downregulated under inductive conditions. Higher levels of *FvTFL1* require longer time to reach the threshold after which the transition to flowering is enabled.

### *FvFT1* is epistatic to *FvTFL1* in control of flowering in woodland strawberry

We previously reported that silencing of *FvFT1* delayed flowering in woodland strawberry *fvtfl1* mutant H4 (Koskela et al. 2012, Rantanen et al. 2014) and reduced the expression of floral meristem identity genes *FvAP1* and *FvLFYa* (Lembinen et al. 2023). Our contrasting findings in this work thus suggest an epistatic interaction between *FvFT1* and *FvTFL1* in the regulation of flowering. To further clarify this, we segregated *FvTFL1* alleles and *FvFT1-*RNAi insertion by generating IT17 × H4 F_2_ lines and observed flowering time in plants grown under different inductive conditions (**Figure 5A**). Under all treatments, *fvtfl1* homozygous mutants (with functional FvFT1) flowered earlier than the *fvtfl1* mutants carrying *FvFT1-*RNAi construct, confirming that FvFT1 activates flowering in the absence of functional FvTFL1. However, the presence of one or two alleles encoding functional FvTFL1 reverted the function of FvFT1. In these cases, silencing of *FvFT1* accelerated flowering in all tested treatments. Furthermore, silencing of *FvFT1* increased the proportion of flowering plants with two functional *FvTFL1* copies under both SD 17°C and LD 11°C treatments. Even under non-inductive LD 17°C conditions, 40% of homozygous and 60% of heterozygous *FvTFL1* plants carrying *FvFT1* silencing construct were able to flower. Moreover, the proportions of flowering plants and the differences in flowering time between homozygous and heterozygous WT *FvTFL1* plants also supported the dosage dependent effect of *FvTFL1* on flowering in woodland strawberry.

**Figure 5.**
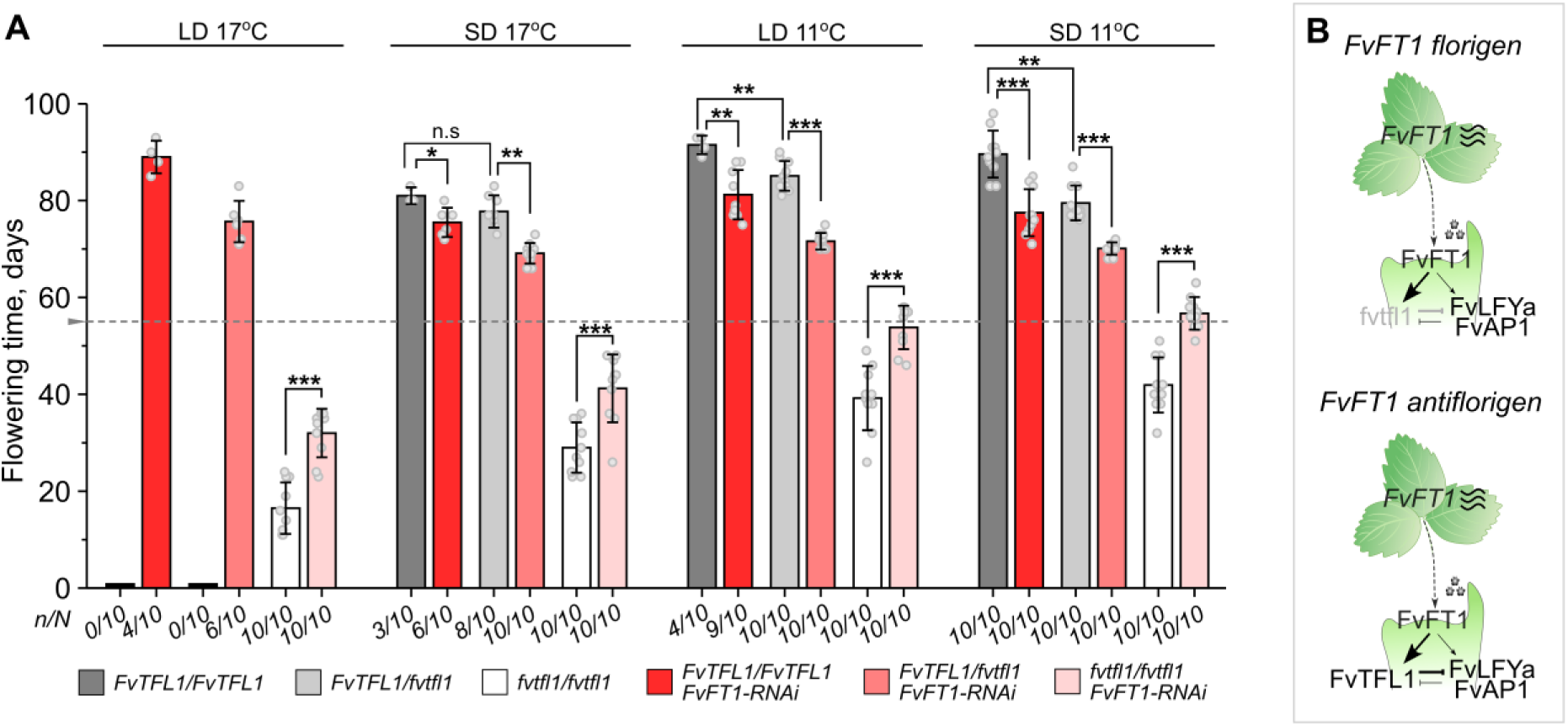
*FvFT1/FvTFL1* epistasis controls flowering in woodland strawberry. **A.** Flowering time of *IT17 × FvFT1- RNAi* F_2_ plants segregating for *FvFT1-RNAi* construct, and *FvTFL1* and *fvtfl1* alleles after eight weeks under the indicated treatments. y-axis shows days since the beginning of the treatments; dashed grey line marks the end of floral inductive treatments after which all the plants were moved to LD 17°C conditions; asterisks indicate the significant differences between selected groups (***, p < 0.001; **, p < 0.01; *, p < 0.05; Tukey’s HSD test); numbers under x-axis denote the number of flowered plants (*n*) and total number of plants in each group (*N*). **B.** The model of flowering time regulation in woodland strawberry.

Altogether our results suggest a model for flowering regulation based on the epistatic interaction between *FvFT1* and *FvTFL1* (**Figure 5B**). When the functional *FvTFL1* allele is present, FvFT1 prevents flowering by promoting *FvTFL1* expression, while in the absence of functional *FvTFL1*, FvFT1 functions as a florigen that activates the expression of *FvLFYa* and *FvAP1* thus promoting flowering. Even in the presence of functional *FvTFL1*, FvFT1 maintains some florigenic activity because its RNAi silencing has a mild downregulating effect on *FvAP1* expression (**Supplemental Figure S8A**), but this activity is masked by *FvTFL1* activation. Moreover, FvFT1 protein does not contain critical amino acid substitutions known to convert FT-like proteins into flowering repressors in other plant species (Hanzawa et al. 2005, Ahn et al. 2006, Pin et al. 2010, Ho & Weigel 2014, Soyk et al. 2017; **Supplemental Figure S8B**). Finally, the expression of *FvFT1* specifically in the leaves (Koskela et al. 2012) is critical for flowering repression mechanism presented here, because ectopic expression of *FvFT1* by the 35S promoter induces flowering even in the presence of *FvTFL1* (**Supplemental Figure S8C**).

## Discussion

In temperate climates, floral transition and preformation of floral buds in autumn/winter is a common life history strategy, yet the patterns of flowering time variation and the underlying molecular mechanisms are understudied. We studied woodland strawberry accessions from diverse bioclimates across Europe and found a geographical pattern of flowering time variation that was shaped primarily by the population structure and winter temperature of the habitat. In concordance with recent report suggesting temperature driven colonization pattern of woodland strawberry from distinct refugia in Europe (Toivainen et al. 2024), earlier flowering accessions typically belonged to the eastern genetic cluster that occupies colder climates, whereas the accessions of the later flowering western cluster were typically found in warmer habitats. Such variation in flowering time is likely attributed to the length of the growing season that creates strong selection pressure on reproductive traits (Austen et al. 2017). The observed stronger correlation between flowering time and winter temperature, rather than latitude, in woodland strawberry, indicates more precise reflection of growing season length by winter temperature, particularly along the longitudinal range. Furthermore, unlike in many other species (Böhlenius et al. 2006, Friedman & Willis 2013, Yang et al. 2019), critical photoperiod seems to play a lesser role in the regulation of floral transition in woodland strawberry (Heide and Sønsteby 2007) and annual growth cycle in Rosaceae fruit trees (Heide and Prestrud 2005).

In habitats with short growing seasons, an early floral transition is crucial to complete flower bud preformation before winter and to advance flowering in the spring, providing more time for fruit development, seed dispersal, and germination (Molau et al. 2005, Schnablová et al. 2020). Also in strawberries, the timing of floral transition in autumn, as observed by meristem microscopy, affects the timing of flowering in the spring (Opstad et al. 2011). Furthermore, in a recent common garden study, southern accessions of woodland strawberry grown in northern Finland showed sporadic and late flowering compared with northern accessions indicating that autumn was too short for floral transition in southern accessions (De-la-Cruz et al. 2025). In our work, the differences in the timing of floral transition between woodland strawberry accessions from colder and warmer climates were evident from the expression patterns of flower meristem identity gene *FvAP1*. We found earlier activation of *FvAP1* in the accessions from cold (northern Norway) than warm (Italy) habitats under flower inductive cool temperatures, indicating a shorter vernalization requirement in the accessions from colder climates. A similar pattern was observed in *A. alpina*. The *A. alpina* accessions from warmer Iberian Peninsula had an obligatory vernalization requirement, whereas in the accessions from colder regions in Alps and Scandinavia this requirement was less common (Wunder et al. 2023). We also found that longer exposure to autumn/winter conditions in the field and controlled climate reduced the overall flowering time variation in our plant collection, while the clinal and genetic cluster related differences remained significant. Strawberry flower buds continue development and reach a more advanced stage when exposed for a longer time to autumn and mild winter temperatures especially in late flowering genotypes (Opstad et al. 2011, Andrés 2025), explaining reduced flowering time variation after longer treatments used in this study. Taken together, variation in the timing of floral transition is a major developmental factor contributing to flowering time variation in woodland strawberry.

At the molecular level, our results indicate that the variation in the expression of woodland strawberry florigen ortholog, *FvFT1*, provides a molecular mechanism controlling flowering time variation. Lower *FvFT1* expression level was associated with colder climate, higher latitudes and earlier flowering, supporting the idea that reduced activity of the photoperiodic pathway may contribute to adaptation to short growing seasons in both annual and perennial plants (Hyun et al., 2019). The geographical patterns in allelic variation of *FT* and contribution of its regulatory elements to flowering time adaptation have been shown in annual Arabidopsis (Strange et al., 2011, Liu et al. 2014). Also in woodland strawberry, we found that differences in the expression of *FvFT1* were associated with the population structure. The plants belonging to the eastern genetic cluster, occupying colder habitats, had lower expression level of *FvFT1* than the plants from the warmer habitats of the western cluster. Major *FvFT1* haplotypes, including highly differentiated SNPs in the regulatory sequences around the gene and in the introns, found in habitats with contrasting winter temperatures in east and west, and their allele specific expression suggested that *cis*-regulatory variation contributes to lower expression of *FvFT1* in the accessions of the eastern genetic cluster. In addition, we found evidence of differential trans-regulation of *FvFT1*. *FvCO*, that is required for the activation of *FvFT1* (Kurokura et al. 2017), and several other putative upstream regulators were differentially expressed together with *FvFT1*, especially in the local populations from northern Norway (Alta and Kåfjord) that pioneered postglacial northward colonization of Europe (Toivainen et al. 2024). Among these genes, *FvCDF2a* and *FvFRI-like* are located within distinct QTL regions controlling early flowering in Kåfjord and Alta, respectively, suggesting convergent evolution of flowering time adaptation in these northern populations with low *FvFT1* expression levels.

Functional analysis confirmed the role of *FvFT1* as the key component of the FvFT1-FvTFL1 flowering repressor module in woodland strawberry. However, in contrast to the previously described FT-like repressors (Pin et al. 2010, Soyk et el. 2017), the flowering repressive property of FvFT1 in strawberry does not arise from critical amino acid substitutions. Instead, the flower repressive function of *FvFT1* is due to an epistatic interaction with *FvTFL1*, encoding a strong floral repressor (Koskela et al. 2012). Upregulation of an antiflorigen in SAM, in response to floral promoting signals, was proposed as a developmental mechanism to protect the SAM from termination in sympodial plants (Tal et al. 2017). In line with this idea, in strawberry, rapid SAM termination occurs when *FvFT1* is overexpressed in *fvtfl1* background, whereas the presence of the functional FvTFL1 prevents termination (Lembinen et al. 2023). Our data shows that such relationship may have further evolved to fine tune the timing of reproductive transition, which terminates the growth axis in sympodial plants. The other woodland strawberry homologs of *FT*, *FvFT2* and *FvFT3* that promote flowering when overexpressed (Gaston et al. 2023), are active in the late inflorescence and flower meristems (Koskela et al. 2017, Lembinen et al. 2023), when *FvTFL1* expression has been suppressed. Given that the formation of inflorescences and flowers takes place in short days of autumn, *FvFT2* and *FvFT3* may compensate for the lack of *FvFT1* in those conditions.

The requirement for vernalization has evolved independently across multiple plant lineages (Yan et al. 2004; Wang et al. 2009; Pin et al. 2010; Bouché et al. 2017), leading to a diversity of adaptive molecular mechanisms that fine-tune the duration and temperature thresholds of vernalization in response to local climates. In winter annual and perennial Brassicaceae, variation in vernalization requirements is primarily governed by differences in the expression levels of *FLC* genes under warm long-day conditions and their downregulation dynamics during vernalization (Albani et al. 2012; Calderwood et al. 2021; Hepworth et al. 2020). In contrast, woodland strawberry, which lacks *FLC* ortholog, uses a different mechanism for flowering time adaptation. FvFT1 enhances the expression of *FvTFL1* under long days and warm temperatures, thus inhibiting flowering. Under short days and cool temperatures, when *FvFT1* expression is repressed, *FvTFL1* is gradually downregulated in the shoot apical meristem enabling activation of inflorescence and flower meristem identity genes (Koskela et al. 2012, Lembinen et al. 2023). Lower *FvFT1* expression level in plants originating from habitats with colder winters contributes to lower initial *FvTFL1* expression level increasing the sensitivity of woodland strawberry to flower inductive conditions across southwest northeast transect, which ultimately contributes to adaptive flowering time variation (**Figure 6**). Such role of *FvTFL1* is widespread in the Rosaceae. According to whole-genome sequencing data, our collection of 202 woodland strawberry accessions exclusively contained functional *FvTFL1* alleles (Toivainen et al. 2024). Furthermore, transcriptional silencing of *TFL1* orthologs is required for flowering in other strawberry species and other species of the Rosaceae family, although the flower inductive signals may vary and repressive mechanisms remain to be shown (Koskela et al. 2012, Iwata et al. 2012, Freiman et al. 2012, Flachowsky et al. 2012, Rantanen et al. 2015, Koskela et al. 2016, Bai et al. 2017, Fan et al. 2023). In conclusion, our results demonstrate how rewiring of a molecular pathway can drive the adaptive variation in flowering time, altogether showing how plants from different lineages yet sharing a similar life-history strategy can utilize different molecular mechanisms to achieve similar adaptive outcome.

**Figure 6.**
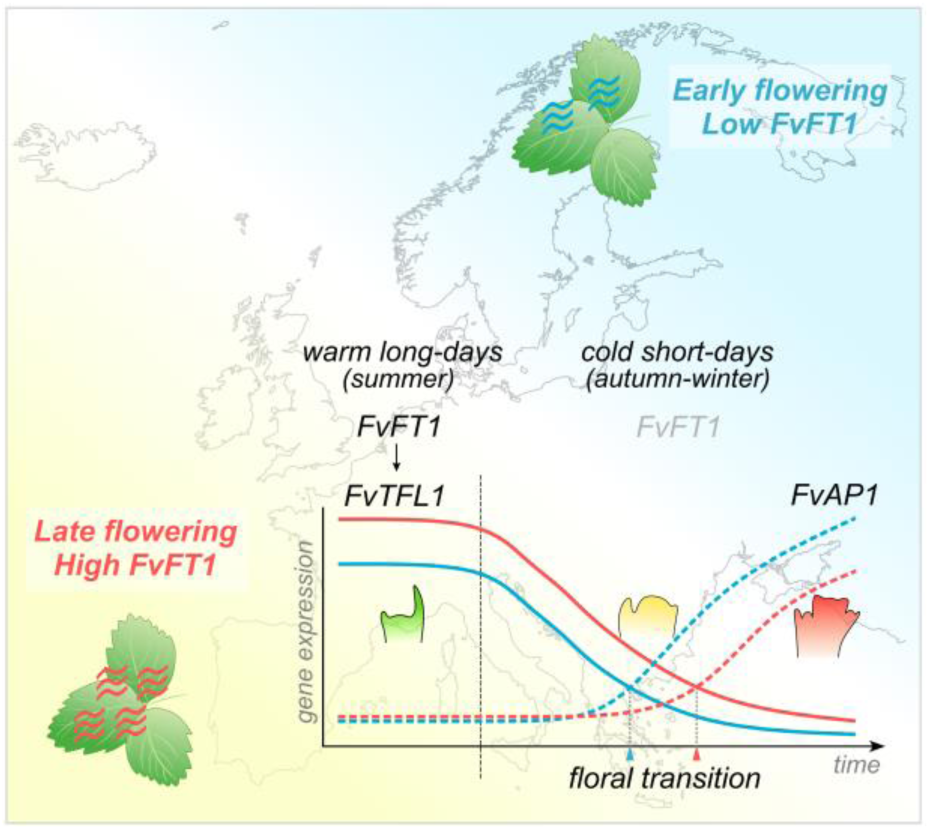
The putative model of flowering time variation in woodland strawberry. Meristem symbols with green, yellow and red color depict vegetative shoot apical meristem, inflorescence meristem and flower meristem, respectively.

## Materials and Methods

### Plant material

Sampling locations and population structure of *F. vesca* accessions are described in Toivainen et al. 2024. *F. vesca fvtfl1* mutant accession *Hawaii-4* (H4) and previously reported *FvFT1* RNA silencing (RNAi) lines in *fvtfl1* (*Hawaii-4*) background were used for gene functional studies (Koskela et al., 2012). To evaluate the effect of *FvFT1* silencing in the presence of functional *FvTFL1*, *F. vesca* accessions FIN56 and IT17 were crossed with H4 (control) and *FvFT1-*RNAi#6 line. To evaluate the epistatic interaction between *FvFT1* and *FvTFL1*, two F_2_ segregating populations were produced by self-pollinating IT17 × H4 F_1_ and IT17 × *FvFT1*-RNAi F_1_ plants. For QTL mapping, two segregating F_2_ populations were produced by self-pollinating IT4 × NOR8 and IT7 × NOR15 F_1_ crosses.

### Growth conditions and phenotyping

Collection of *F. vesca* accessions, QTL mapping F_2_ populations and transgenic plants were maintained in a glasshouse at 18°C temperature and under 18/6 h (day/night) (LD) photoperiod. The plants were illuminated with 150 µmol m^-2^ s^-1^ light intensity, using high-pressure sodium lamps (Airam 400W, Kerava, Finland). Plants were irrigated with tap water supplemented with fertilizers (Ferticare; N-P-K 7-4-27) and Calcinit (YARA; N 15, Ca 19). For the experimental treatments, plants were clonally propagated by rooting stolons to peat pellets (Jiffy growing solutions), while attached to mother plants. After rooting, plants were potted to 10 x 10 cm pots and pre-grown for 4–6 weeks before starting floral inductive treatments.

In the accession collection, floral transition was induced in a closed glasshouse with controlled climate or under ambient temperature and photoperiod in an open glasshouse (rooftop without walls). In controlled climate experiments, plants were grown for six weeks under indicated temperature and photoperiod conditions (SD - 12/12 h; LD – 18/6 h day/night). After SD 16°C and LD 11°C treatments plants were moved to LD 20°C to observe flowering. In SD 11°C/6°C/-2°C experiment, in addition to six weeks in SD 11°C, plants were further exposed to SD 6°C for six weeks and then placed to cold storage room (−2°C, 0/24 h day/night) for two months and moved to LD 20°C to observe flowering. In field experiments, plants were exposed to ambient photoperiod and temperature (Helsinki, Finland) from mid-August until October 12^th^ (2017, Field Aug-Oct) or until January 4^th^ (2018, Field Aug-Jan) before moving to the greenhouse (LD 20°C) for flowering observations. In Field Sep-Jun experiment, plants were exposed to ambient temperature and photoperiod until December 1^st^ (2021) and moved to cold storage room (−2°C, 0/24 h day/night) until April 12^th^ (2022); after that the plants were moved back to open glass house for flowering observations.

For QTL mapping, flowering was induced in F_2_ populations in the greenhouse at SD 17°C for six weeks, after that plants were moved to LD 18°C room for flowering observation. For the gene functional and expression analyses, the selected accessions, F_1_ hybrids, and F_2_ segregants were grown in walk-in growth rooms. Plants were illuminated with 140 µmol m^-2^ s^-1^ light intensity (AP67, Valoya, Finland). Clonally propagated plants were pre-grown at 17°C and under 18h/6h day/night photoperiod and then subjected to four different conditions: 18/6 h photoperiod at 17°C (control), 12/12 h photoperiod at 17°C, 18/6 h photoperiod at 11°C and 12/12 h photoperiod at 11°C. WT accessions and F_1_ hybrids were kept under the treatments for six weeks, F_2_ populations for eight weeks. Plants were irrigated with tap water supplemented with fertilizer (N-P-K 13-7-20, Puutarhan Kesä, GreenCare). The number of replicates for each experiment is reported in figures or figure legends. After the treatments plants were moved to the greenhouse (18/6 h, 18°C) and flowering time was recorded as the date of the first open flower.

### RNA extraction and gene expression analyses

For gene expression analyses, leaf discs and shoot apices were used. Leaf discs of 0.5 x 0.5 cm were sampled at ZT4. A single leaf disc was placed into an Eppendorf tube representing one biological replicate. Shoot apices containing the apical meristem were dissected from the main growth axis. Two to three apices were pooled into one Eppendorf tube, representing one biological replicate. Samples were immediately frozen in liquid nitrogen and stored at −80°C. Total RNA was extracted from leaf and apex samples as in Mouhu et al. (2009) and treated with rDNase (Macherey-Nagel GmbH, Düren, Germany).

For qRT-PCR, 500ng of total RNA was used for cDNA synthesis using ProtoScriptII reverse transcriptase (NEB, US). qRT-PCR was performed using a LightCycler 480 SYBR Green I Master kit (Roche Diagnostics, Indianapolis, US) in a Roche LightCycler qPCR machine (Roche Diagnostics, Indianapolis, US) with three technical replicates for each of the tested genes. *FvMSI1 (FvH4_7g08380)* was used as a stable reference gene (Mouhu et al. 2009). The qRT-PCR primers used in this study are listed in **Supplemental Table S4**.

The expression of *FvAP1* and *FvTFL1* in the apices of Norwegian and Italian accessions (**Figure 4D – E**) was analyzed using Nanosting assay (NanoString Technologies, Seattle, USA) at the Institute of Biotechnology (Helsinki, Finland). RNA samples were diluted to 20ng/µl. The NanoString probes were designed and synthesized by NanoString (**Supplemental Table S4**). Samples were normalized using geometric mean of *FvMSI1* (*FvH4_7g08380*) and *FvACTIN* (*FvH4_7g22410*) as reference genes.

### RNA sequencing and co-expression analysis

For bulk RNA sequencing, total RNA was isolated from leaf tissues collected four hours after dawn (ZT4) from three biological replicates from 12 accession (six western and six eastern) grown under 18/6 h (day/night) photoperiod at 18°C in the greenhouse. RNA sequencing libraries were prepared using TruSeq Total RNA kit (Illumina) after removal of ribosomal RNA with Pan-Plant riboPOOL-probes (siTOOLs Biotech) and MagSi-STA 1.0 (Magtivio) beads. The sequencing of 100 bp fragments was carried out on the Illumina NextSeq500 sequencer. Raw fastq reads were quality-trimmed using Trimmomatic v0.39 (Bolger et al. 2014) in single-end mode with the following parameters: SLIDINGWINDOW:5:20, LEADING:5, TRAILING:5, and MINLEN:50. Cleaned reads were aligned to the *Fragaria vesca* reference genome using STAR v2.7.10a (Dobin et al. 2013), with the genome indexed using the v4.0.a2 annotation provided by Genome Database for Rosaceae website. Only uniquely mapped reads were retained for downstream analysis. Gene-level quantification was performed using HTSeq-count v0.13.5 (Anders et al. 2015) in unstranded mode.

Co-expression analysis was performed on bulk RNA sequencing data from 12 accessions. Prior to co-expression analysis, normalization and filtering was performed using edgeR package pipeline (Robinson et al. 2010). Gene-level CPM (counts per million) values from 36 RNA-seq libraries were obtained using normalized library sizes. Only genes with sufficient counts were retained as determined using `filterByExpr` function. Based on multidimensional scaling (MDS) plot and sample clustering, three samples were removed from further analyses (**Supplemental Figure S6A - B**). Differentially expressed genes between accessions from eastern and western genetic clusters were estimated using edgeR pipeline (fold change > 2, FDR < 0.05, p-value < 0.05). Co-expressed genes were identified using weighted correlation network analysis package (R/WGCNA, Langfelder and Horvath 2008). Soft-thresholding power for network construction was estimated as 12. Minimum size of the clusters was set to 50. Threshold for merging clusters (`mergeCutHeight`) was set to 0.25. Other parameters were kept as default. Pearson’s correlation was estimated between module (cluster) eigengene values and traits (e.g. flowering time, bio11, latitude). For visualization, CPM values were converted to FPKMs.

### Allele-specific RNA sequencing

Total RNA was isolated from leaf tissues collected at ZT4 from three biological replicates from IT7 × NOR15 F1 plants. Allele Specific RNA sequencing was carried out with Illumina Novaseq SE sequencer to produce 100 + 100 bp paired-end reads. The quality of the sequencing data was confirmed with FastQC and trimmed with Trimmomatic (Bolger et al. 2014). The sequences were mapped to *F. vesca* genome v4.0.a2 with STAR 2-pass mode (Dobin et al. 2013). Read counts from mapping were checked with HTSeq (Anders et al. 2015) using htseq-count. The GATK best practices were followed for variant calling (DePristo et al. 2011). and for GATK tools (McKenna et al. 2010) version 4.2.6.6 was used. The SAM files from mapping were sorted and duplicates were marked with Picard (https://broadinstitute.github.io/picard/). Then the bam file was processed with GATK SplitNCigarReads and read groups were assigned with Picard. The variants were called with HaplotypeCaller (Poplin et al. 2017) and GATK SelectVariant was used to filter biallelic SNPs from the output VCF file. GATK ASEReadCounter was used to count the allele-specific counts for biallelic SNPs. The Chi-Square test was used to estimate which SNPs had significant p-values for allelic imbalance.

### *FvFT1* sequence analyses

Genetic differentiation (F_ST_, Weir & Cockerham 1984) was calculated for all SNPs and accessions of the eastern (N = 111) and western (N = 88) genetic clusters (Toivainen et al. 2024; **Supplemental Data S5**). Top 5% SNPs with the highest F_ST_ values (F_ST_ ≥ 0.409357) were then identified followed by searching such SNPs in a 10 kb region of *FvFT1* gene. A closer look at the haplotypes of the most highly differentiated SNPs indicated that Icelandic accessions despite generally belonging to the western genetic cluster (Toivainen et al. 2024) contained the SNPs typical for the accessions of the eastern cluster, suggesting a recent introgression in that region. We recalculated F_ST_ excluding the Icelandic accessions (N = 18; **Supplemental Data S6**), identified the top 1% SNPs with the highest F_ST_ values (F_ST_ ≥ 0.592048) across the genome, and identified *FvFT1* haplotypes.

FT and TFL1 protein sequences of Arabidopsis, tomato, sugar beet and woodland strawberry were retrieved from Phytozome v13 database (**Supplemental Data S7**). Protein sequences were aligned using Clustal Omega web-based software (Madeira et al. 2024) with default parameters.

### Genotyping

QTL populations were genotyped through whole genome sequencing of individual F2 lines. Library preparation and sequencing was performed as in Toivainen et al. (2024), except Novaseq S1 system was used for sequencing 100 + 100 bp paired-end reads (**Supplemental Data S8**). Mapping and SNP calling for the QTL-mapping populations were conducted with a similar protocol as previously applied to the 202 accessions (Toivainen et al. 2024). Briefly, Cutadapt (Martin 2017) trimmed fastq reads were mapped against *Fragaria vesca* reference genome (Edger et al. 2018) with BWA-MEM (Li 2013). Duplications were marked with the Picard tool MarkDuplicates. GATK v 3.8. HaplotypeCaller-function was used to call variants in single samples, and then, variants were jointly called across all samples using the GenotypeGVCFs-function (**Supplemental Data S9**).

A 2-bp deletion in *fvtfl1* sequence in H4 WT and transgenic (*FvFT1-RNAi*) crosses was genotyped using fragment analysis (ABI3730xl, ThermoFisher Scientific) with FAM labeled primers (Koskela et al. 2012, **Supplemental Table S4**) at the University of Helsinki. The peaks were analyzed using *Peak Scanner*™ Software Version *1.0* (Applied Biosystems).

### QTL Mapping

Linkage map was constructed using Lep-MAP3 (Rastas, 2017) from the variants of the vcf file. The family structure was validated using the IBD module (using a random subset of 0.1% of the markers in the vcf), which calculates the identity-by-descent (IBD) values between all the individuals. These values were >=0.5 only between IT7 × NOR15 (F_2_) and NOR8 × IT4 (F_2_) families and joining these high-IBD individuals clustered individuals into these two families. The two selfing families were given to Lep-MAP3 as two full-sib families by adding two dummy parents to both families to obtain a pedigree accepted by Lep-MAP3. The “parental” genotypes were obtained by ParentCall2 module. Filtering2 was used to filter too distorted markers (non-default parameter `dataTolerance=0.0001`). Then the remaining markers were clustered into 7 linkage groups with `SeparateChromosomes2` (first using `lodLimit=12`, then `lodLimit=13` and `map=map_lodLimit12`), and finally, `OrderMarkers2` (`selfingPhase=1`, to phase the data according to selfing cross) was used to obtain a linkage map. The map positions were extracted from the number of crossovers output by Lep-MAP3, separately for both populations.

The genome assembly quality was visually verified by constructing a scatter plot of linkage and genome assembly positions of the markers. The genome and the map positions were visually consistent (**Supplemental Figure S9A and B**) and 3 extra contigs could be added to the assembled chromosomes. The exact positions for these contigs were decided based on gaps in the scaffolds and the contig-contig alignments constructed using HaploMerger2 software (Huang et al, 2017). The final linkage map was constructed by evaluating the map in the refined genome assembly order (`OrderMarkers` with parameter `evaluateOrder`). This reduced noise in the map, especially from the map ends. The parental genotypes were phased as well from the parental phase information output by Lep-MAP3’s `OrderMarkers2`. The data for QTL mapping was constructed with `map2genotypes.awk` script on the final map and the number of markers was reduced by keeping only the first and the last marker of each unique map position.

QTL mapping was performed using *R/qtl2* packgage (Broman et al., 2019). 89 individuals from IT7 × NOR15 (F_2_) and 92 individuals NOR8 × IT4 (F_2_) populations were used for mapping (**Supplemental Data S10**). Genotype probabilities were estimated with `error_prob=0.002` to calculate kinship matrix. A linear mixed model genome scan was performed using leave-one-chromosome-out method. LOD score thresholds were estimated by permutation analysis (`perm = 10000`). All estimated thresholds were below four. Candidate genes within estimated confidence intervals were searched using the genome annotation v4.0.a2 provided by Genome Database for Rosaceae.

### Statistical analyses

Bioclimate data for 19 bioclimatic variables and mean monthly temperatures of plant collection sites were extracted from the WorldClim 2.0 dataset at 2.5-minute spatial resolution based on sample coordinates (**Supplemental Data S1**). Pearson’s correlation coefficients between flowering time, bioclimate variables and plant location were calculated. P-values were estimated with Bonferroni correction. Non-linear relationship between flowering time, bio11 and latitude was tested using generalized additive modelling (GAM) with penalized cubic regression spline (`bs=“cs”`) implemented in *R/mgcv* packgage (Wood 2004). Spline parameter k was set at 5 to avoid overfitting. An additional mixed model (GAMM) using experimental treatment as a mixed effect was fitted to all data to estimate average smoothers (**Figure 1C – D**). The effect of the genetic clusters on flowering time responses to bio11 and latitude was analyzed using linear regression (Bayesian). Data from each experiment were analyzed separately. Posterior distributions and confidence intervals of regression parameters were estimated using *R/INLA* package (Lindgren & Rue, 2009). The flowering time data from common garden experiment, F_1_ hybrids, selected NOR and IT accessions and F_2_ segregants were analyzed using ANOVA followed by pairwise t-tests with Tukey’s HSD. All analyses were performed in R version 4.3.0 (R Core Team 2023) and are summarized in **Supplemental Table S5**.

## Acknowledgments

The research was funded by the Research Council of Finland (grant nr 317306 and 327409 to T.H.) and through the 2019-2020 BiodivERsA joint call for research proposals, under the BiodivClim ERA-Net COFUND programme, and with the funding organization Research Council of Finland (grant nr 344726 to T.H.). S.L. was funded by a personal grant from Finnish Cultural Foundation (grant nr 00221168). G.F. and Q. Z. were supported by grants from China Scholarship Council (grant nr 201706510014 to G.F. and 202007960007 to Q.Z.). J.A. was funded by Doctoral Programme of Plant Sciences (University of Helsinki, Finland). The study included material from the “Professor Staudt Collection” maintained in Dresden, Germany at the Hansabred GmbH & Co. KG and from the IFAPA *Fragaria* collection funded by IFAPA Project PR.CRF.CRF202200.002 of the European Agricultural Fund for Rural Development, within the Rural Development Program of Andalusia 2014-2022. We thank Eija Takala and Marjo Kilpinen for laboratory assistance and Katriina Palm for maintaining the plant materials. The analysis of high-throughput sequencing data was performed using CSC - IT Center of Science-high-performance computing cluster.

