## Supplementary Figures for "*FvFT1-FvTFL1* epistasis drives flowering time adaptation in woodland strawberry"

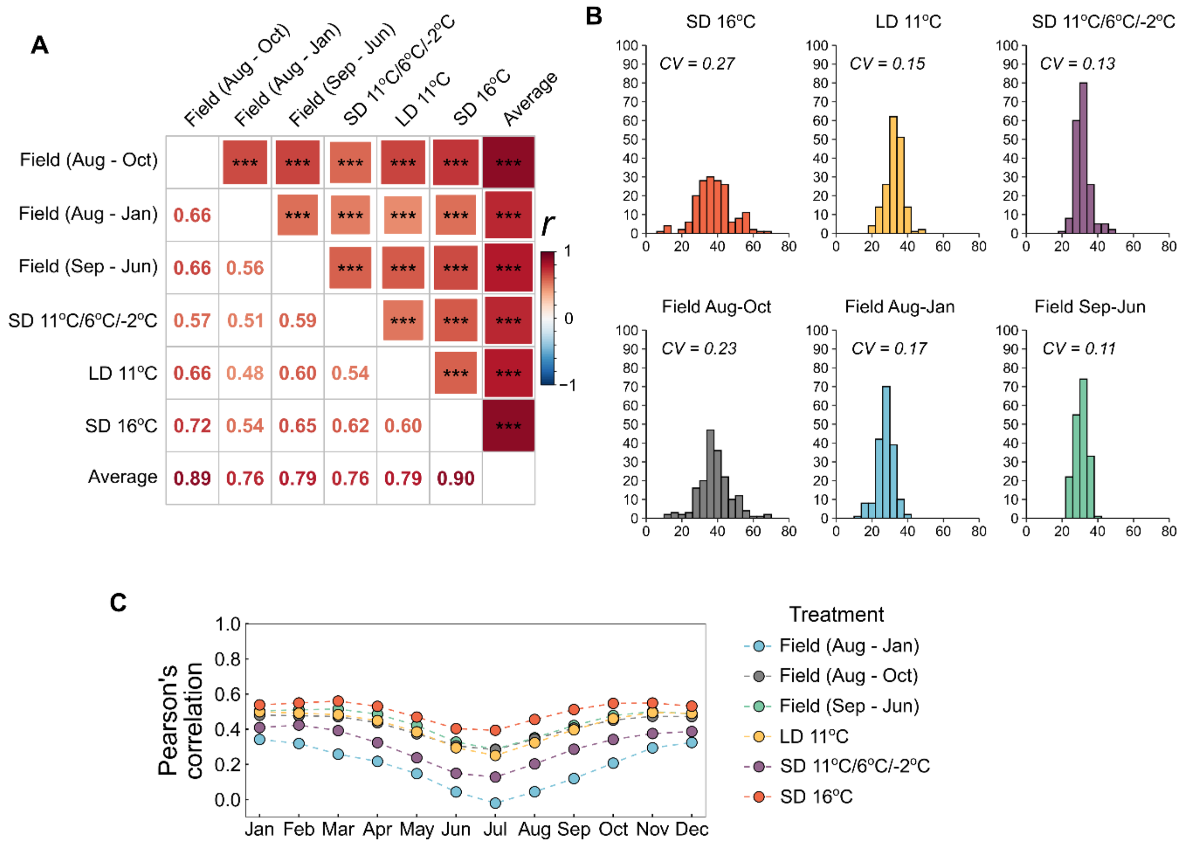


**Supplemental Figure S1. A.** Pearson's correlation between flowering time of woodland strawberry accessions after different experimental treatments. Colors of the square indicate correlation strength, asterisks indicate significant correlations (***, p ≤ 0.0001; **, p ≤ 0.001; *, p ≤ 0.01, Bonferroni adjusted). **B.** Histograms showing flowering time variation after different experimental treatments. CV – coefficient of variation. **C.** Pearsons correlations between flowering time and mean monthly temperatures. Colors indicate experimental treatments.

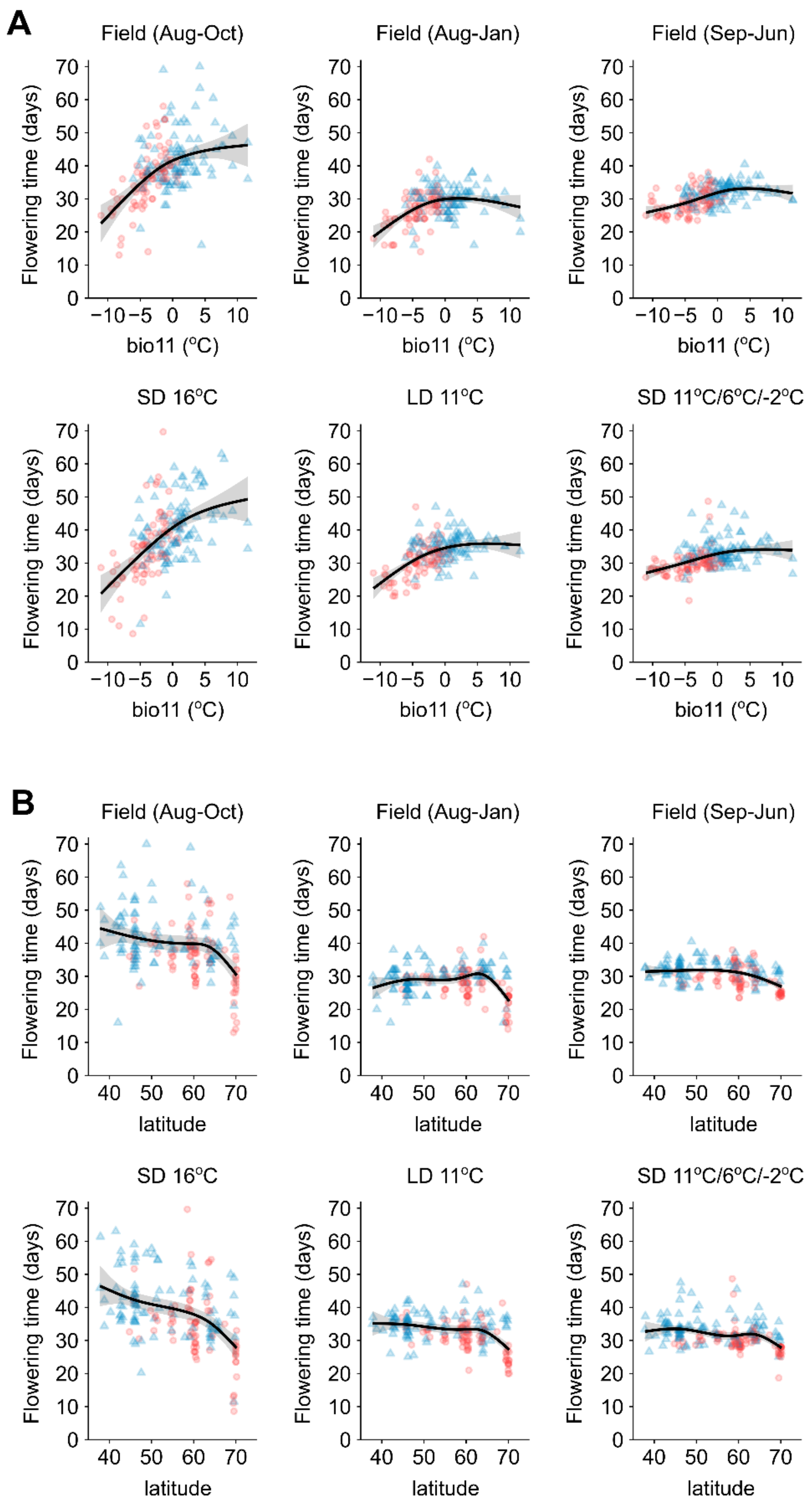


**Supplemental Figure S2.** Non-linear patterns in flowering time along bio11 (A) and latitude (B) of original habitats of woodland strawberry accessions from individual experiments. Color of points denotes genetic cluster of the accession (western – blue, eastern – red). Black curves with shaded areas show fitted GAM models with confidence intervals (95%).

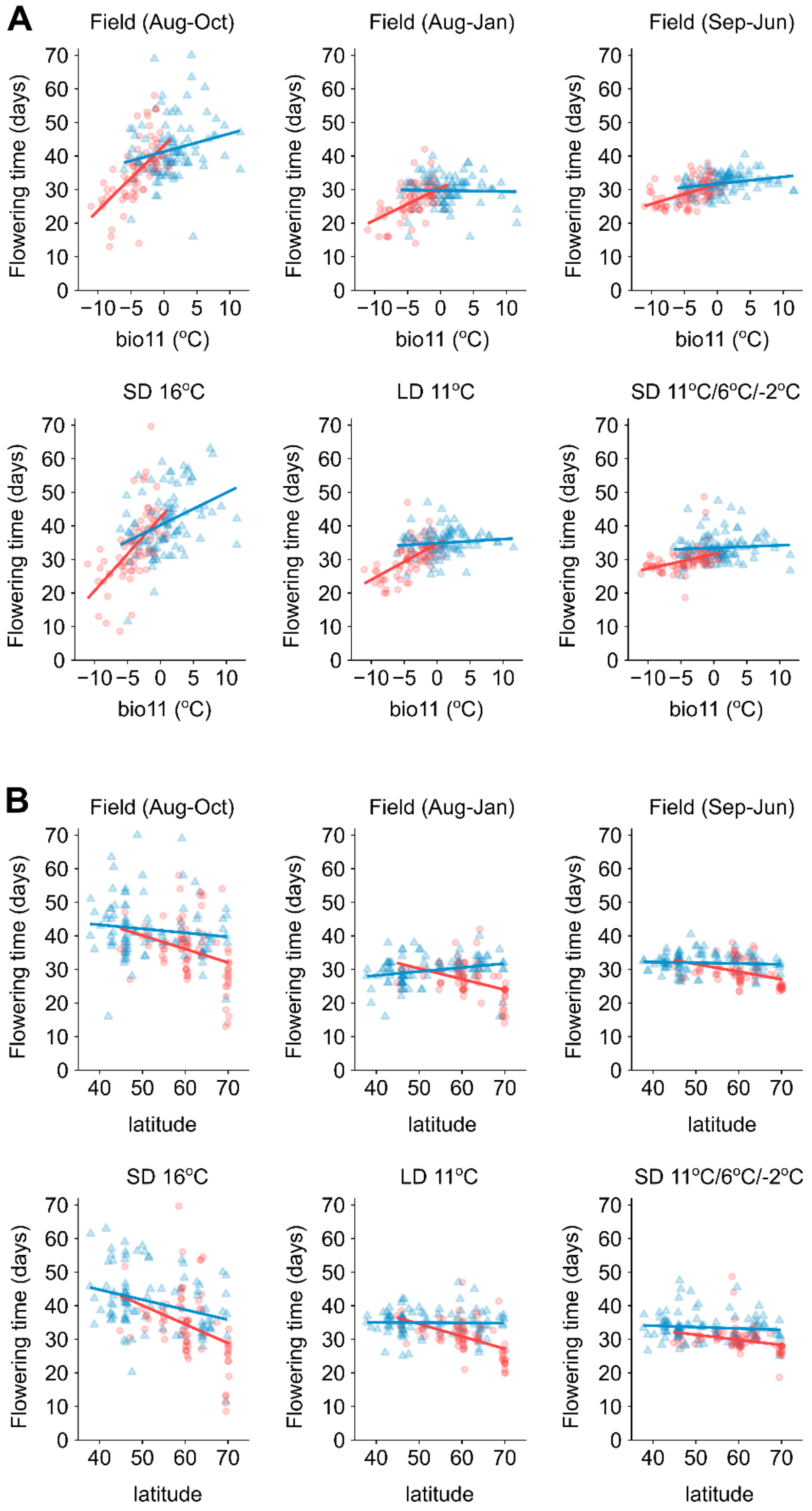


**Supplemental Figure S3.** Linear regression models showing differences in flowering time responses to bio11 (A) and latitude (B) between eastern (red) and western (blue) genetic clusters of woodland strawberry in individual experiments.

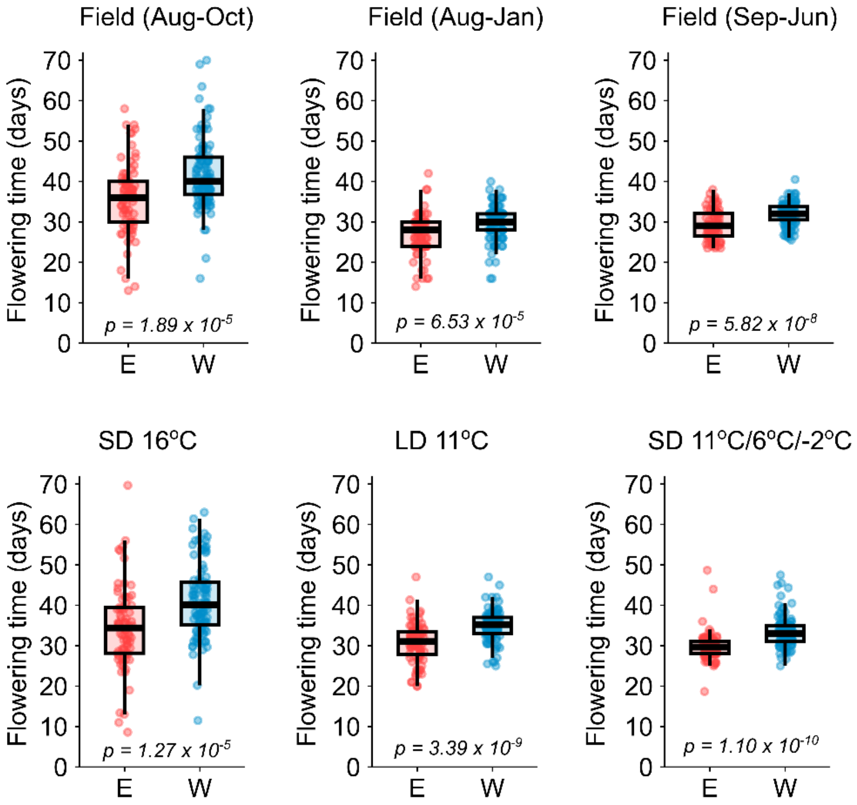


**Supplemental Figure S4.** Comparison of flowering time between the woodland strawberry accessions of the eastern (E) and western (W) genetic clusters in indicated experiments (t-test).

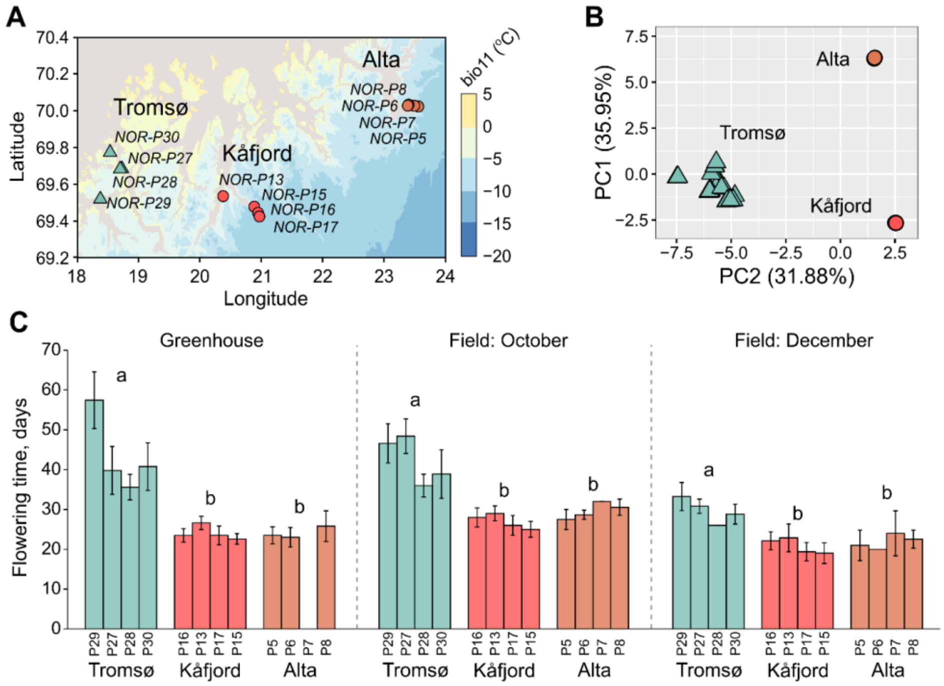


**Supplemental Figure S5.** Flowering time variation in woodland strawberry populations from northern Norway. **A.** Locations of sampled populations from three areas in northern Norway. **B.** PCA plot showing structure of northern Norwegian populations. Symbols indicate genetic cluster (circles - eastern, triangles - western) according to Toivainen et al. (2024). **C.** Average flowering time of woodland strawberry populations after flower induction in a greenhouse (12-h SDs at 11°C) or in a common garden (Helsinki). Letters indicate significant differences between areas ($p < 0.05$, GLMM (Poisson) with Tukey's HSD); y-axis shows days to flowering after the end of the treatments.

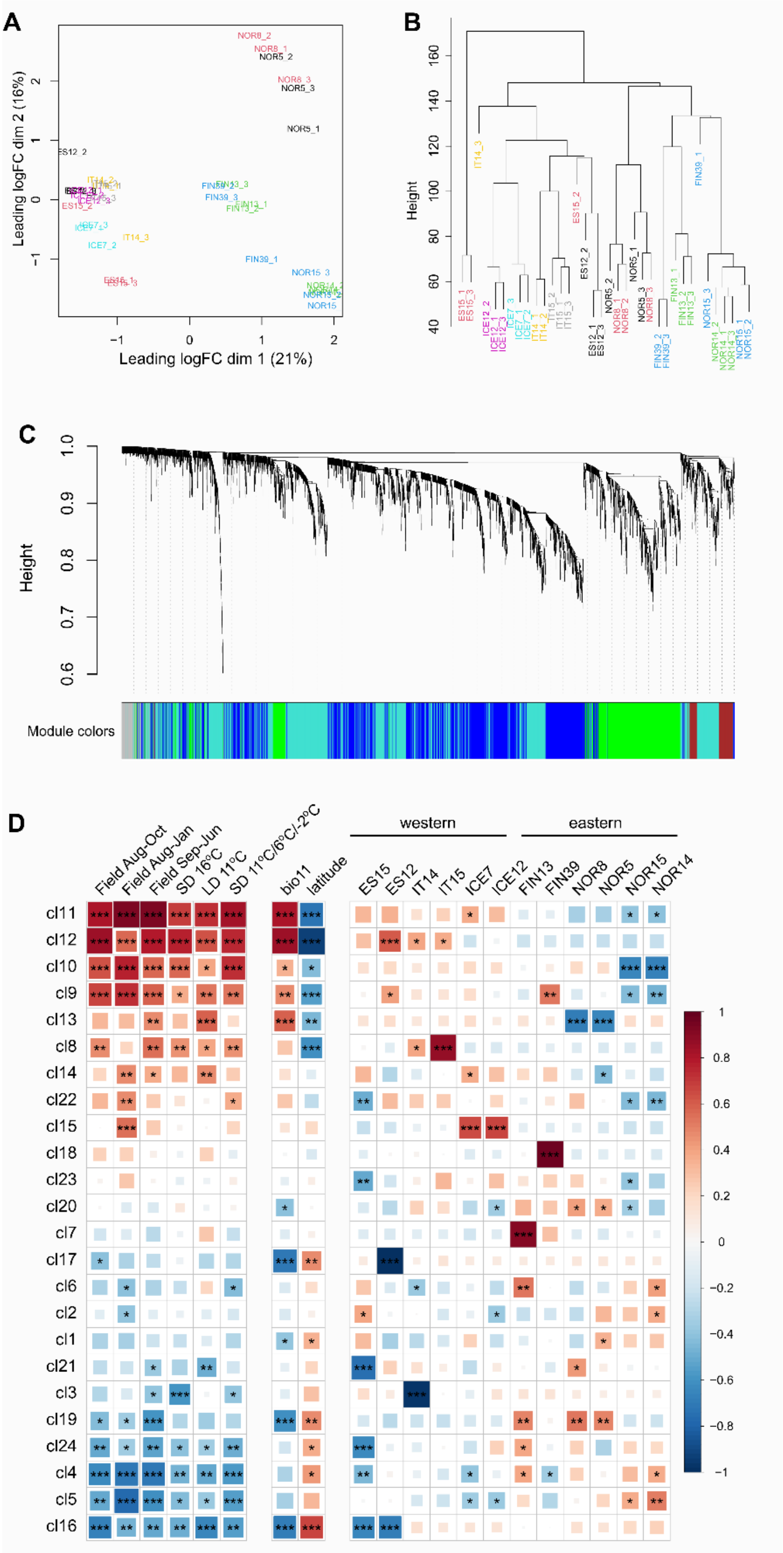


**Supplemental Figure S6.** Summary of the RNAseq analysis of the leaf samples collected under 18-h long days from 12 woodland strawberry accessions. **A - B.** Multidimensional scaling (MDS) plot (A) and a dendrogram (B) showing clustering of RNAseq samples. **C.** WGCNA dendrogram showing gene clustering. **D.** Extended heatmap showing correlations between RNAseq clusters and phenotypes.

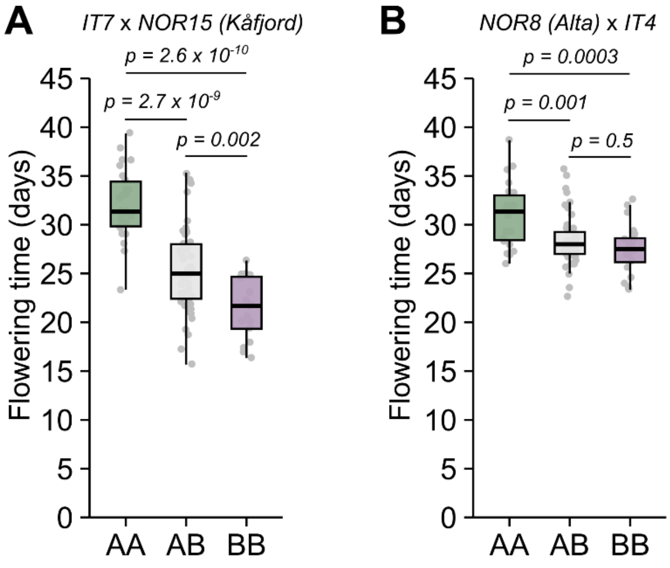


**Supplemental Figure S7.** QTL effects of the highest significance markers (Tukey's HSD) in the indicated woodland strawberry $F_2$ mapping populations. Y-axis shows days to flowering after the end of the 6-week treatment under 12-h short days at 17°C.

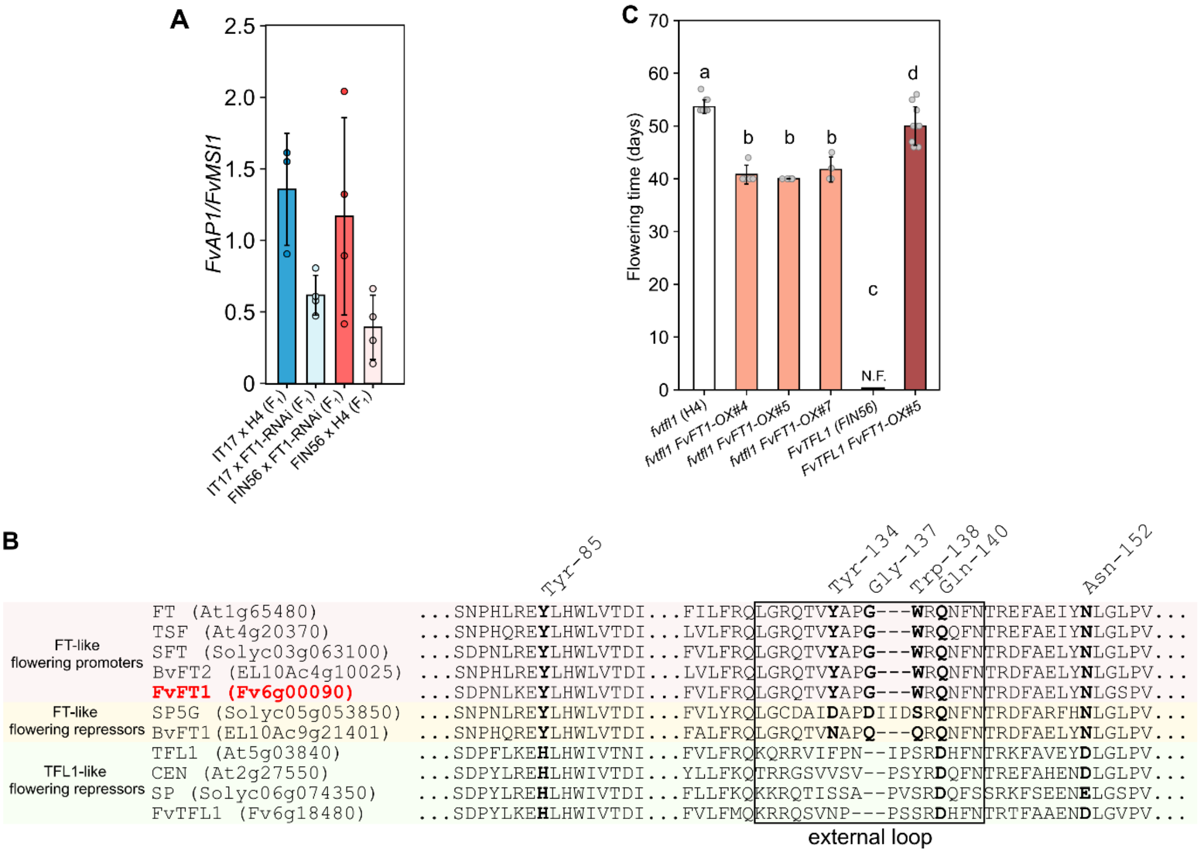


**Supplemental Figure S8. Woodland strawberry *FvFT1* function as a flowering promoter. A.** A zoom in on **Figure 4B** showing expression of *FvAP1* before the start of a floral inductive treatment (week 0; LD (18h/6h day/night) and 17°C) in control and *FvFT1-RNAi* crosses. **B.** Alignment of FT and TFL1 proteins from Arabidopsis, beet, tomato and woodland strawberry. Critical amino acids conferring flowering promoter or repressor functions are highlighted in bold. **C.** Flowering time (days after potting) in *FvFT1* overexpression lines (CaMV 35S) in different *FvTFL1* backgrounds under long-days (18h/6h day/night) in 17°C. Different letters indicate statistically significant differences (p-value < 0.05, Tukey's HSD).

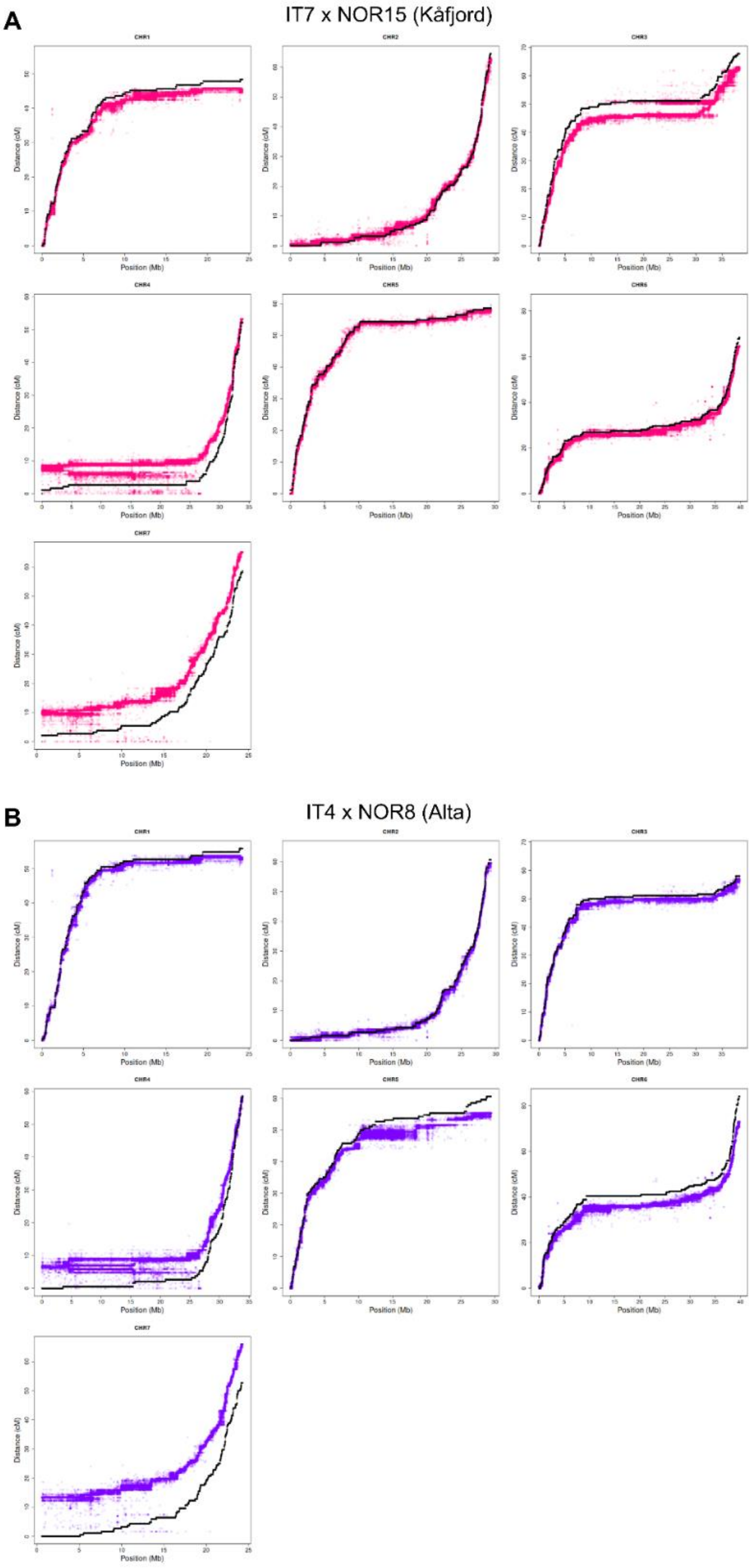


**Supplemental Figure S9.** Marey maps for all seven linkage groups (chromosomes) of indicated woodland strawberry $F_2$ mapping populations. The black points display the map with markers in the genome assembly order, whereas the purple and red points show the map constructed independent of the genome.
